# Interferon-γ signaling synergizes with LRRK2 in human neurons and microglia

**DOI:** 10.1101/2020.01.30.925222

**Authors:** Silvia De Cicco, Dina Ivanyuk, Wadood Haq, Vasiliki Panagiotakopoulou, Aleksandra Arsić, David Schöndorf, Cong Yu, Maria-Jose Perez, Ruggiero Pio Cassatella, Meike Jakobi, Nicole Schneiderhan-Marra, Ivana Nikić-Spiegel, Thomas Gasser, Michela Deleidi

## Abstract

Increasing evidence suggests a role for interferons (IFNs) in neurodegeneration. Parkinson’s disease (PD) associated kinase LRRK2 has been implicated in IFN type II (IFN) response in infections and nigral neuronal loss. However, whether and how LRRK2 synergizes with IFN-γ still remains unclear. Here, we employed dopaminergic (DA) neurons and microglia differentiated from patient induced pluripotent stem cells to unravel the role of IFN-γ in LRRK2-PD. We show that IFN-γ induces LRRK2 expression in both DA neurons and microglial cells. LRRK2-G2019S, the most common PD-associated mutation, sensitizes DA neurons to IFN-γ by decreasing AKT phosphorylation. IFN-γ suppresses NFAT activity in both neurons and microglia and synergistically enhances LRRK2-induced defects of NFAT activation. Furthermore, LRRK2-G2019S negatively regulates NFAT via calcium and microtubule dynamics. Importantly, we uncover functional consequences of the reduction of NFAT activity in both cell types, namely defects of neurite elongation and alteration of microglial activation profile and motility. We propose that synergistic IFN-γ/LRRK2 activation serves as a direct link between inflammation and neurodegeneration in PD.

## Introduction

Emerging data suggest that immune dysfunction contributes, not only to the progression, but also to the onset of neurodegenerative diseases including Parkinson’s disease (PD) ^1^. In this respect, human genetics and functional genomics studies indicate that interferon-mediated signaling pathways, both type I and type II, contribute to brain ageing and human neurodegenerative diseases ^2, 3, 4^. With regard to PD, there is evidence for a link between expression network signatures of disease loci and IFN-γ signaling ^5^. Recently, IFN-γ response has also been linked to the selective vulnerability of dopaminergic (DA) neurons ^6^. IFN-γ is a cytokine primarily produced by T-lymphocytes and natural killer cells as part of the immune response against pathogens ^7^. IFN-γ plays a key role in both the innate and adaptive immune responses and contributes to macrophage activation, up-regulation of major histocompatibility complex (MHC), and modulation of T helper response ^8^. Besides its role in immune responses against pathogens, IFN-γ is also fundamental to normal brain physiology ^9^. The high incidence of postencephalitic parkinsonism following the pandemic influenza towards the end of World War I, had already suggested a selective vulnerability of DA neurons to pathogen-driven immune responses. Indeed, related pathological studies revealed the presence of inflammatory encephalitis in the midbrain and basal ganglia, without central nervous system viral invasion ^10^. Supporting a possible role of IFN-γ in the disease process, increased levels of IFN-γ in the serum, as well as increased production of IFN-γ by peripheral CD4+ T cells, have been detected in PD patients compared to healthy controls ^11, 12^. Moreover, α-synuclein and IFN-γ has been found within the significant co-expression of α substantia nigra of PD patients ^13^. Experimental models also support this connection, whereby IFN-γ has been linked to selective, progressive, DA neurodegeneration in rodents ^11, 14, 15^. More recent studies also point towards the interaction between PD-related genes and IFN-γ induced DA neuronal loss ^16^. The study by Kozina et al. shows that human pathogenic mutations in the leucine rich repeat kinase 2 (*LRRK2*) gene, the most common genetic cause of familial and sporadic PD ^17^, synergizes with lipopolysaccharide (LPS)-induced inflammation to potentiate DA neurodegeneration through IFN-γ-mediated immune responses ^16^. Interestingly, *LRRK2* is an IFN-γ target gene ^18, 19^ and its expression is increased in immune cells upon IFN-γ stimulation and exposure to pathogens ^18, 20, 21, 22, 23^. Genetic polymorphisms in *LRRK2* have also been associated with Crohn’s disease (CD) and leprosy ^24, 25, 26^, suggesting overlapping pathogenetic mechanisms among chronic inflammatory diseases, infections and PD. LRRK2-PD manifests similar clinical phenotypes to idiopathic PD, displaying a strong age-dependent development of PD symptoms.

Notably, PD patients carrying the *LRRK2* G2019S mutation have distinct peripheral inflammatory profiles ^27^. Despite the evidence of a link between IFN type II immune response and PD, mechanisms through which IFN-γ contributes to neurodegeneration and interacts with PD genes, such as *LRRK2*, are still unknown. Based on these premises, we aimed to investigate mechanisms of IFN-γ-mediated neurotoxicity and to examine the role of the most common pathogenic missense mutation, LRRK2 G2019S, in modulating IFN-γ signaling.

## Results

### IFN-γ induces LRRK2 expression in human neurons

To examine the impact of IFN type II signaling on human neurons, we employed induced pluripotent stem cell (iPSC)-derived neural precursor cells (NPCs) Control NPCs were differentiated into neuronal cultures enriched in DA neurons and treated with IFN-γ for 24 hrs. Since previous work has shown that IFN-γ induces LRRK2 in immune cells ^18, 20, 21, 22^, we examined LRRK2 mRNA and protein levels in untreated and IFN-γ treated neuronal cultures. IFN-γ stimulation significantly increased LRRK2 γ mRNA and protein levels in human neurons (Fig. 1A, B). To evaluate whether LRRK2 missense mutations influence the IFN-γ-driven induction of LRRK2, we used NPC lines derived from patients carrying the LRRK2 G2019S mutation and corresponding isogenic controls ^28, 29^. As previously reported ^29^, isogenic NPCs were efficiently differentiated into DA neurons without significant differences among patients and controls (Supplementary Fig. 1A). LRRK2 protein expression was similar among different lines (Supplementary Fig. 1B). IFN-γ treatment also increased LRRK2 protein levels in LRRK2 G2019S PD patient-derived neurons (Fig. 1C). IFN-γ treatment did not affect neuronal cell viability, as assessed by LDH assay (Supplementary Fig. 1C). To examine the specificity of the effects observed with IFN-γ, control neurons were treated with 200 IU/mL IFN-γ, 100 ng/mL LPS or 100 IU/mL γ IL-1β for 24 hrs and LRRK2 mRNA levels were measured using qRT-PCR. Only IFN-γ efficiently induced LRRK2 expression in human neurons (Fig. 1D).

**Figure 1.**
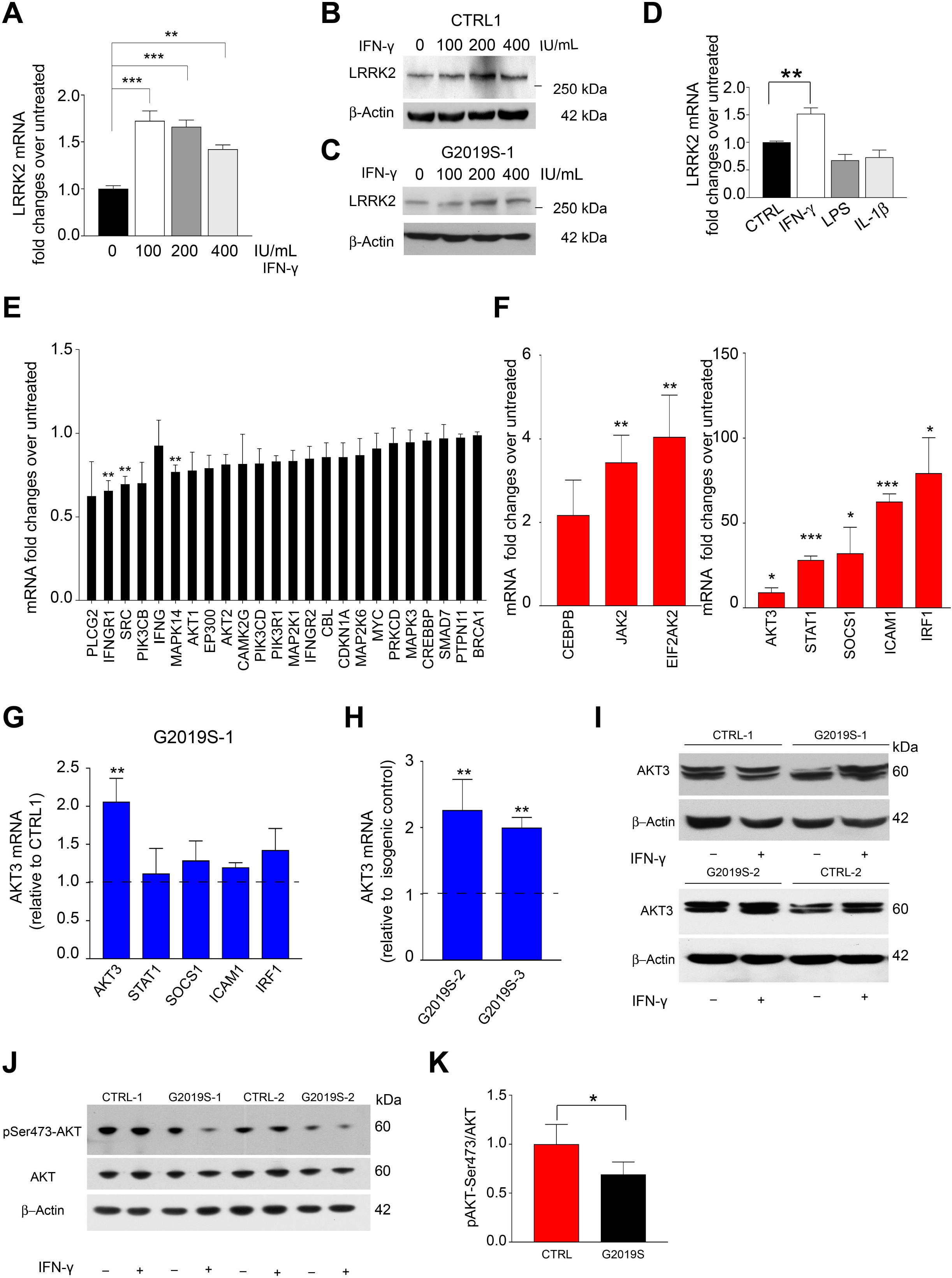
IFN-γ induces LRRK2 in human iPSC-derived neurons and decreases AKT phosphorylation in LRRK2 G2019S neurons. **(A-D)** IFN-γ treatment induces LRRK2 levels in human iPSC-derived neurons. (A) LRRK2 mRNA levels in control iPSC-derived neurons treated with IFN-γ for 24 hrs at the indicated concentrations. LRRK2 mRNA levels, measured by qRT-PCR, are expressed as fold changes of untreated (mean ± SEM, one-way ANOVA, Bonferroni post hoc, ***P<0.001, **P <0.01; n=3 independent experiments). (B, C) Representative western blot for LRRK2 expression in control (B) and isogenic LRRK2 G2019S (C) iPSC-derived neurons treated with IFN-γ at the indicated concentrations. (D) LRRK2 mRNA levels in control iPSC-derived neurons treated with 200 IU/mL IFN-γ, 100 ng/mL LPS, or 100 IU/mL IL-1β for 24 hrs (mean ± SEM, one-way ANOVA, Bonferroni post hoc, **P<0.01; n=3 independent experiments). **(E, F)** IFN-γ transcriptional signature genes in control iPSC-derived neurons treated with 200 IU/mL IFN-γ for 24 hrs measured by qRT-PCR and expressed as fold changes over untreated. Down-regulated (E) and upregulated (F) genes are shown (mean ± SEM, t-test treated vs untreated, ***P<0.001, **P<0.01, *P<0.05; n=3 independent experiments). **(G)** mRNA levels of AKT3, STAT1, SOCS1, ICAM1, IRF1 in LRRK2 G2019S (G2019S-1) neurons treated with 200 IU/mL IFN-γ, expressed as fold changes over corresponding IFN-μ treated isogenic control (CTRL-1) (mean ± SEM, t-test, **P<0.01; n= 3 independent experiments). **(H)** mRNA levels of AKT3 in LRRK2 G2019S neurons (G2019S-2, G2019S-3) treated with 200 IU/mL IFN-γ, expressed as fold changes over corresponding treated isogenic controls (CTRL-2, CTRL-3) (mean ± SEM, t-test, **P<0.01; n=3 independent experiments). **(I)** Representative Western blot of AKT3 in LRRK2 G2019S neurons (G2019S-1, G2019S-2) and corresponding isogenic controls (CTRL-1, CTRL-2). Treatment with 200 IU/mL IFN-γ is indicated. **(J)** Representative Western blot of p-AKT Ser473 in LRRK2 G2019S neurons (G2019S-1, G2019S-2) and corresponding isogenic controls (CTRL-1, CTRL-2). Treatment with 200 IU/mL IFN-γ is indicated. **(K)** Quantification of pAKT/AKT level in LRRK2 G2019S neurons and isogenic controls. Data are expressed as fold changes of IFN-γ treated over untreated and normalized to CTRL (mean ± SD, t-test, *P<0.05; n= 4 independent experiments).

### LRRK2 G2019S alters IFN-γ signaling in human neurons

To characterize IFN-γ response in human neurons, healthy control NPC lines were differentiated into DA neurons and treated with 200 IU/mL IFN-γ for 24 hrs. IFN-γ-regulated genes, selected according to the INTERFEROME gene expression database (Interferome DB v2.0) ^30^, were profiled by qRT-PCR in vehicle- and IFN-γ-treated neurons. Interferon Gamma Receptor 1 (*IFNgR1*), SRC proto-oncogene, non-receptor tyrosine kinase (*SRC*), and mitogen-activated protein kinase 14 (*MAPK14*) were significantly down-regulated (Fig. 1E), whereas Janus kinase 2 (*JAK2*), eukaryotic translation initiation factor 2 alpha kinase 2 (*EIF2AK2*), AKT serine/threonine kinase 3 (*AKT3*), signal transducer and activator of transcription 1 (*STAT1*), suppressor of cytokine signaling 1 (*SOCS1*), intercellular adhesion molecule 1 (*ICAM1*), and interferon regulatory factor 1 (*IRF1*) were significantly up-regulated in IFN-γ treated compared to vehicle treated neurons (Fig. 1F). To examine whether LRRK2 G2019S mutation influences neuronal IFN-γ response, we examined the expression of the most significant IFN-γ upregulated genes in LRRK2 G2019S mutant NPC-derived neurons and corresponding isogenic controls. Among the upregulated genes analysed, we found significantly higher mRNA levels of AKT3 in LRRK2 G2019S mutant neurons compared to the corresponding isogenic controls upon IFN-γ stimulation (Fig. 1G). These results were confirmed in two different pairs of LRRK2 G2019S patient neurons and isogenic gene corrected controls (Fig. 1H). At the protein level, we detected a modest non-significant increase of AKT3 in IFN-γ-treated cells (Fig. 1I). We then examined whether LRRK2 G2019S influences AKT phosphorylation state. Given the lack of phospho-AKT3-specific antibodies, an antibody recognizing all AKT isoforms was used. IFN-γ treatment reduced AKT phosphorylation in both LRRK2 G2019S and control neurons (Fig. 1J); however, AKT phosphorylation was significantly reduced in LRRK2 G2019S compared to control neurons upon IFN-γ treatment (Fig 1J, K).

### LRRK2 G2019S reduces nuclear NFAT3 in human neurons

Next, we investigated the possible impact of reduced AKT phosphorylation and reasoned that this could have a negative effect on the translocation and transcriptional activity of the nuclear factor of activated T-cells (NFAT) ^31, 32^. First, we employed HEK293 NFAT reporter cell lines. We transfected them with LRRK2 wild-type (wt) or LRRK2 G2019S constructs (Fig. 2A), and measured NFAT transcriptional activity in response to phorbol-12-myristate-13-acetate (PMA) and ionomycin. In line with previous reports ^33^, we found that overexpression of LRRK2 wt inhibits NFAT-dependent transcriptional activity (Fig. 2B). Moreover, NFAT activity was decreased to a greater extent in cells overexpressing LRRK2 G2019S (Fig. 2B). Next, HEK293 NFAT reporter cells (overexpressing LRRK2 wt or LRRK2 G2019S) were treated with IFN-γ and stimulated with PMA/ionomycin. IFN-γ treatment decreased NFAT activity and the most potent effect was observed in patient LRRK2 G2019S overexpressing cells (Fig. 2C). Next, we sought to investigate the link between IFN-γ/LRRK2 and NFAT in DA neurons. We first examined the expression levels of the Ca^2+^ dependent NFAT isoforms (NFAT1, NFAT2, NFAT3, and NFAT4) in human iPSC-derived neurons and microglia by qRT-PCR (Supplementary Fig. 1D). As NFAT3 exhibited the highest expression in neurons, we focused on this family member. To further elucidate the link between LRRK2 and NFAT in PD, we transfected LRRK2 G2019S PD neurons and corresponding isogenic controls with an NFAT3-GFP reporter plasmid and examined NFAT localization by quantitative immunofluorescent analysis. Consistent with the luciferase assay data, we observed decreased nuclear NFAT localization in patient LRRK2 G2019S neurons compared to isogenic controls (Fig. 2D, E). Interestingly, IFN-γ treatment further decreased the nuclear localization of NFAT3 and the most potent effect was observed in patient LRRK2 G2019S neurons (Fig. 2D, F). To confirm that LRRK2 acts downstream of IFN-γ, we generated LRRK2 knockout (LRRK2 KO) iPSCs (Supplementary Fig. 1E). LRRK2 KO iPSCs and isogenic controls were differentiated into neurons and treated with IFN-γ or vehicle. We observed increased nuclear NFAT localization in LRRK2 KO neurons compared to isogenic controls, whereas IFN-γ did not have a significant impact on NFAT localization in LRRK2 KO neurons (Fig. 2D, G). We then performed Western blot analysis for NFAT levels in LRRK2 G2019S PD patient neurons and controls with and without IFN-γ treatment. No changes were observed in total NFAT levels at basal conditions, whereas a significant reduction was observed upon treatment with IFN-γ in both controls and LRRK2 G2019S PD patient neurons (Supplementary Fig. 1F). Similar results were obtained with nuclear and cytosolic fractionation experiments (Supplementary Fig. 1G, H). Notably, we observed a reduction of NFAT levels upon IFN-γ treatment (Supplementary Fig. 1F-H). Given that IFN-γ activates the proteasome ^34, 35^, we examined whether the reduction of total NFAT levels observed in IFN-γ-treated neurons could be mediated by proteasome activation. To this end, control neurons were treated with IFN-γ with or without proteasome inhibitor, MG132, and total levels of NFAT were assessed by Western blot. Interestingly, proteasome inhibition restored NFAT levels in IFN-γ treated neurons (Supplementary Fig. 11). Taken together, these data indicate that IFN-γ leads to NFAT degradation by the proteasome and that the presence of the mutation G2019S enhances the retention of NFAT in the cytosol by reducing AKT phosphorylation.

**Figure 2.**
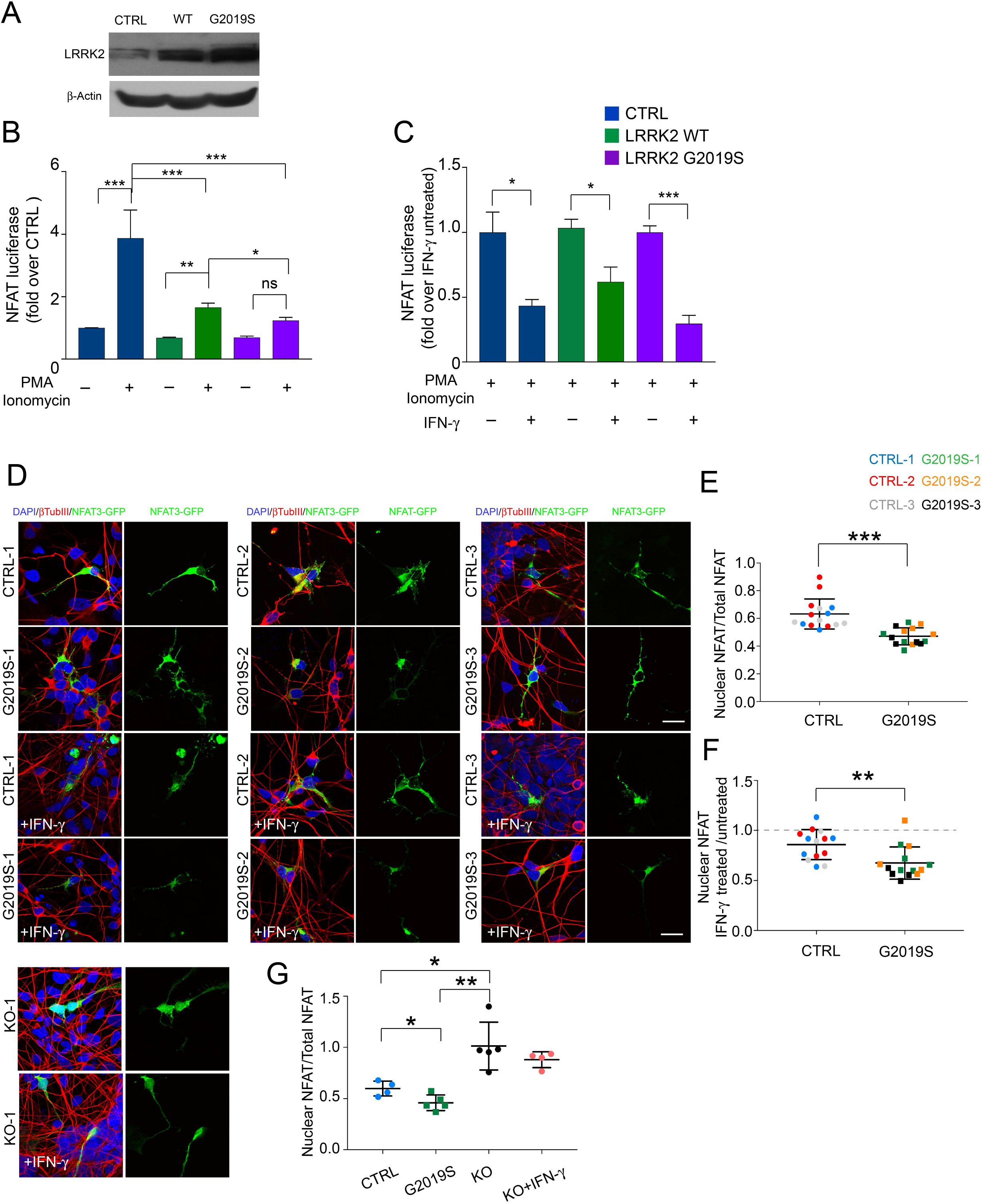
LRRK2 inhibits nuclear NFAT3 shuttling in HEK293 and human neurons. **(A)** Western blot for LRRK2 in HEK cells transfected with wt or LRRK2 G2019S. **(B)** NFAT luciferase assay in HEK293 NFAT reporter cells transfected with wt or LRRK2 G2019S and stimulated with 1 μM ionomycin and 40 ng/mL PMA for 24 hrs (mean ± SD, one-way ANOVA, Bonferroni post hoc, ***P<0.001, **P<0.01, *P<0.05; n=3 independent experiments). **(C)** NFAT luciferase assay in HEK293 NFAT reporter cells transfected with LRRK2 wt or LRRK2 G2019S and stimulated with PMA/Ionomycin for 24 hrs with or without 200 IU/mL IFN-γ. Data are expressed as fold changes of IFN-γ treated over corresponding PMA/Ionomycin stimulated cells (mean ± SD, t-test, ***P<0.001, *P<0.05; n=3 independent experiments). **(D)** Representative confocal microscopy images of LRRK2 G2019S, KO and corresponding control neurons transfected with vector encoding NFAT3-GFP (green) treated with 200 IU/mL IFN-γ for 24 hrs or left untreated; blue, DAPI staining; red, βTubIII. **(E)** Quantification of NFAT3-GFP subcellular localization expressed as nuclear NFAT3-GFP fluorescent intensity normalized on total cell body intensity (mean ± SD, t-test, ***P<0.001; n=5 independent experiments). **(F)** Quantification of NFAT3-GFP subcellular localization expressed as nuclear NFAT3-GFP fluorescent intensity normalized on total cell body intensity. Data are expressed as ratio of IFN-γ treated over corresponding untreated (mean ± SD, t-test, **P<0.01; n=5 independent experiments). **(G)** Quantification of NFAT3-GFP subcellular localization expressed as nuclear NFAT3-GFP fluorescent intensity normalized on total cell body intensity (mean ± SD, t-test, *P<0.05, **P<0.01; n=5 independent experiments).

### LRRK2 alters the ER-related neuronal calcium response to IFN-γ

Since NFAT1-4 family members are regulated by Ca^2+^ signaling, we examined LRRK2 G2019S-related changes in Ca^2+^ homeostasis. Fura-2 live-cell Ca^2+^ imaging was performed in two isogenic pairs and revealed that LRRK2 G2019S PD neurons have higher cytosolic Ca^2+^ basal levels compared to isogenic controls (Fig. 3A, B). Upon KCl depolarization, LRRK2 G2019S PD neurons showed significantly reduced Ca^2+^ influx and longer recovery time compared to controls (Fig. 3A, B), suggesting a defect in the intracellular buffering capacity. Upon Sarco-Endoplasmic Reticulum Ca2+-ATPase (SERCA) inhibition with thapsigargin (TPH), we observed a significantly reduced release of Ca^2+^ from the endoplasmic reticulum (ER) of mutant cell lines compared to controls, indicating a reduced capacity of the ER to uptake and store Ca^2+^ in LRRK2 G2019S mutant neurons (Fig. 3C). To examine the effect of chronic IFN-γ treatment on neuronal Ca^2+^ dynamics, LRRK2 G2019S mutant neurons and controls were treated with 200 IU/mL IFN-γ for 24 hrs and the ER Ca^2+^-release in response to TPH was assessed by live-cell Ca^2+^ imaging. We observed a significant reduction of the ER Ca^2+^-release induced by TPH upon IFN-γ-treatment in control lines (Fig. 3D). On the other hand, no significant difference was observed in IFN-γ-treated LRRK2 G2019S mutant neurons, confirming a pre-existing depletion of ER Ca^2+^-stores (Fig. 3D).

**Figure 3.**
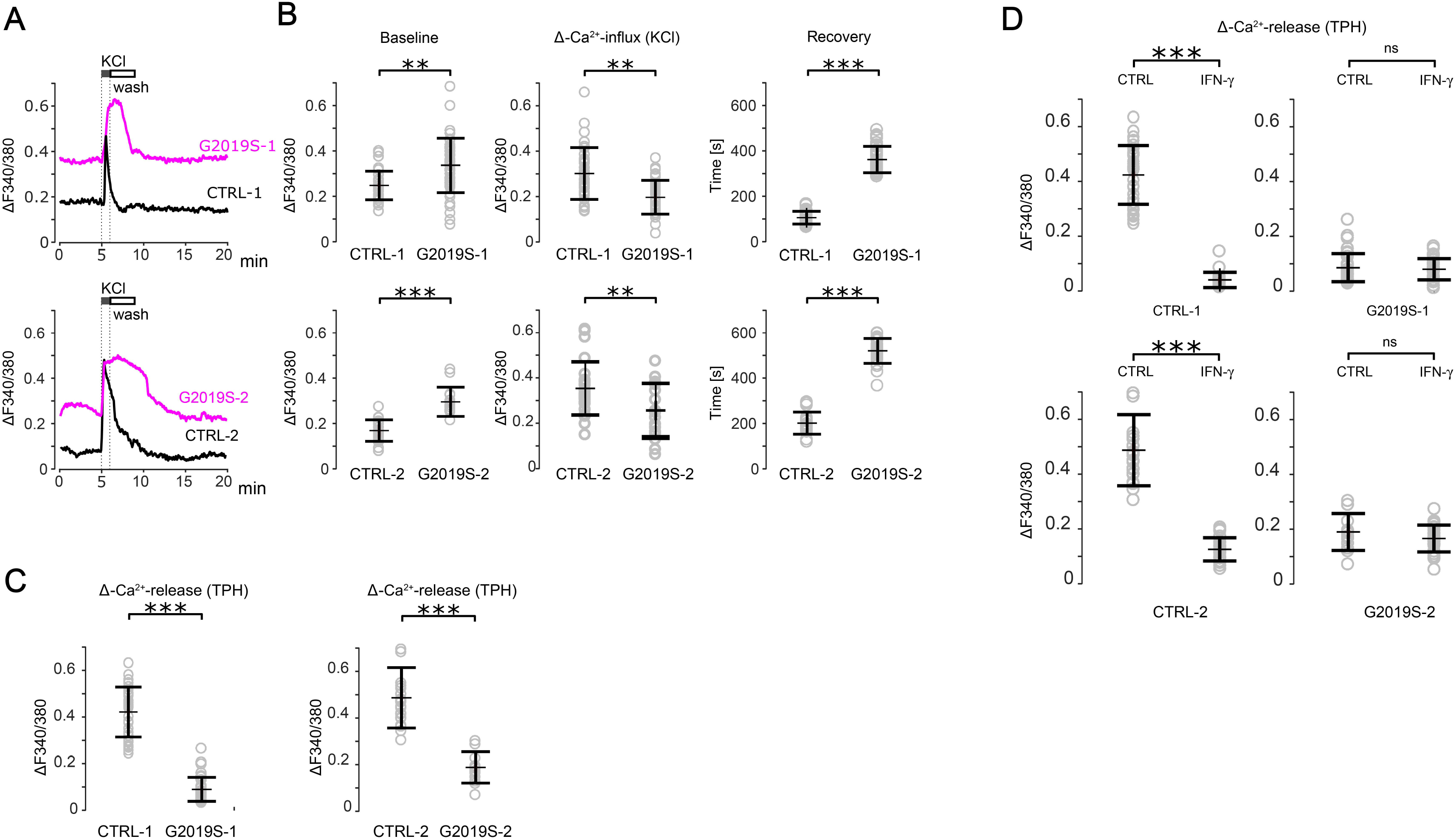
LRRK2 G2019S alters intracellular Ca^2+^ dynamics and Ca^2+^ response to IFN-γ. Intracellular Ca^2+^ dynamics were measured after KCl stimulation in human iPSC-derived neurons (CTRL-1, CTRL-2, G2019S-1, and G2019S-2) using fura-2 AM ratiometric acquisition. **(A)** Representative Ca^2+^ imaging of single neuron traces is shown (normalized data). KCl indicates time of given stimulus. **(B)** Intracellular Ca^2+^ levels at baseline (left panel), Ca^2+^ influx upon KCl stimulation (middle panel), and recovery time after stimulation (right panel) are shown (mean ± SD, t-test, ***P<0.001, **P<0.01; n=3 independent experiments, at least 10-15 single cells per each cell line per experiment were analysed). **(C)** ER Ca^2+^ release upon thapsigargin (TPH) treatment (500 nM, 15 min) in LRRK2 G2019S and isogenic control neurons. Quantification of Ca^2+^ peak amplitude (Δ-Ca^2+^ release) is shown (mean ± SD, t-test, ***P<0.001; n=3 independent experiments, 10-15 cells per experiment were analyzed). **(D)** Quantification of TPH mediated Δ-Ca^2+^ release in isogenic LRRK2 G2019S and control neurons pretreated with 200 IU/mL IFN-γ for 24 hrs or left untreated (mean ± SD, t-test, ***P<0.001; n=3 independent experiments).

### LRRK2 regulates NFAT shuttling through microtubules

Integrity of the microtubule network is necessary for NFAT translocation and T-cell activation ^36^. Furthermore, LRRK2 interacts with and regulates several cytoskeletal components in a variety of cell types. LRRK2 is part of the NRON [noncoding (RNA) repressor of NFAT] complex that also contains IQGAP1 ^33, 38^, a scaffold protein that, among other functions, regulates cytoskeleton architecture, including actin filaments and microtubules, and coordinates their interactions during cell migration at leading edges. To examine whether IQGAP1 is involved in LRRK2 mediated defects of NFAT shuttling, we knocked down IQGAP1 in HEK293 NFAT reporter cell lines (Fig. 4A, B). Knockdown of IQGAP1 increased NFAT activity, both at basal condition and upon PMA/Ionomycin stimulation (Fig. 4C, D). Interestingly, IFN-γ treatment reduced NFAT activity in control but not in IQGAP1 KD cells, suggesting a role for IQGAP1 in IFN-γ mediated negative modulation of NFAT shuttling (Fig. 4E). Furthermore, IQGAP1 KD partially rescued the reduction of NFAT activity in LRRK2 wt and LRRK2 G2019S overexpressing cells, suggesting a role for IQGAP1 in LRRK2-mediated changes on NFAT translocation (Fig. 4F,G).

**Figure 4.**
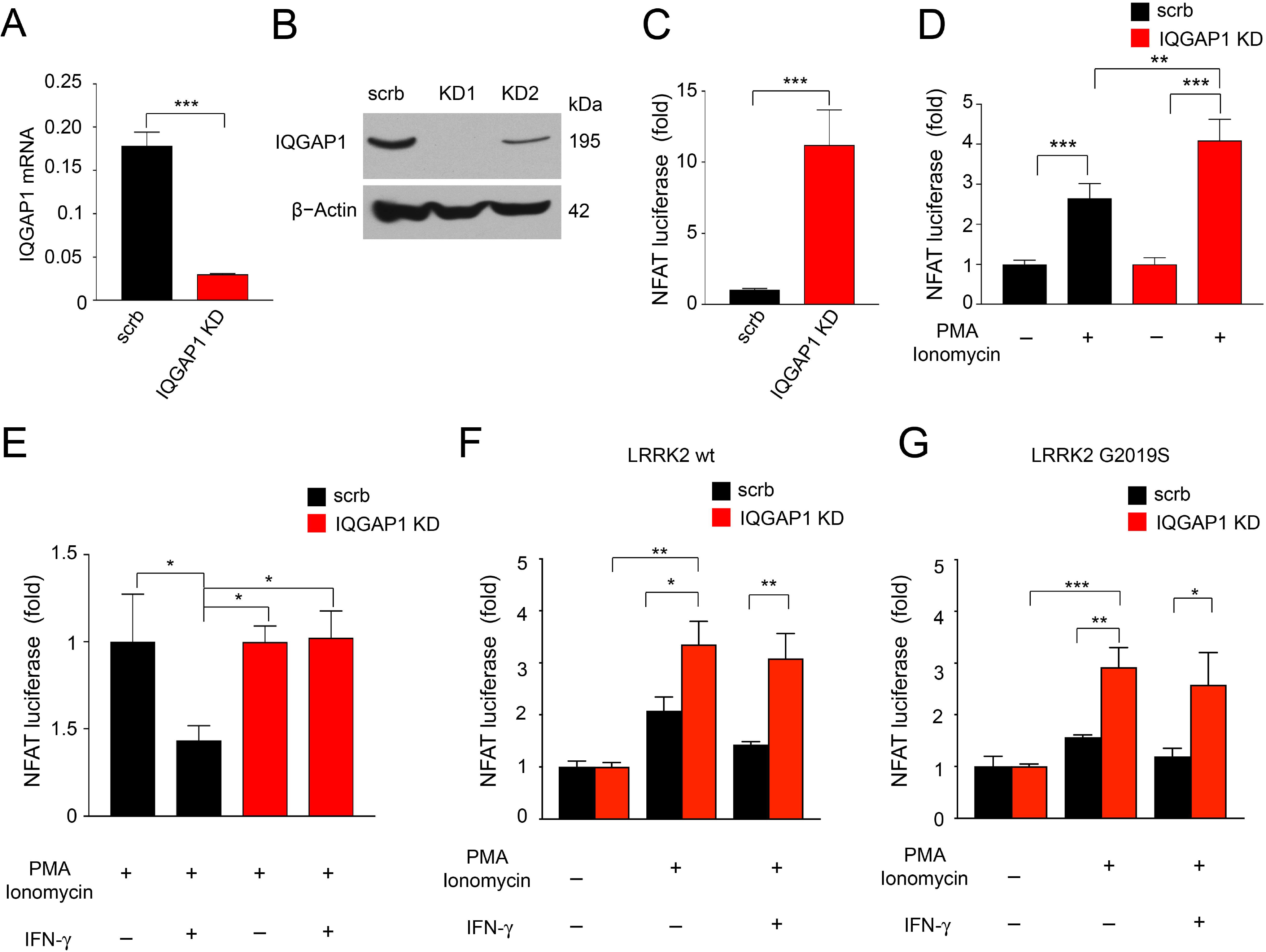
LRRK2 regulates NFAT shuttling through microtubules. **(A)** Knockdown efficiency of IQGAP1 in HEK293 NFAT reporter cells determined by qRT-PCR and normalized to scramble (scrb) non-targeting shRNA (mean ± SD, t-test, ***P<0.001, n=3). **(B)** Representative Western blot for IQGAP1, showing knockdown efficiency. **(C)** NFAT luciferase assay in scrb and IQGAP1 KD HEK293 NFAT reporter cells at basal level. Data are expressed as fold change over scrb (mean ± SD, t-test, ***P<0.001, n=3 independent experiments). **(D)** NFAT luciferase assay in scrb and IQGAP1 KD HEK293 NFAT reporter cells with or without PMA/Ionomycin stimulation. Data are expressed as fold change over corresponding vehicle treated control (mean ± SD, one-way ANOVA, Bonferroni post hoc, ***P<0.001, **P<0.01; n=3 independent experiments). **(E)** NFAT luciferase assay in scrb and IQGAP1 KD HEK293 NFAT reporter cells stimulated with PMA/Ionomycin for 24 hrs, and 200 IU/mL IFN-γ for 24 hrs, where indicated. Data are expressed as fold changes over corresponding PMA/Ionomycin treated control (mean ± SD, one-way ANOVA, Bonferroni post hoc, *P<0.05; n=3). **(F)** NFAT luciferase assay in scrb and IQGAP1 KD HEK293 NFAT reporter cells transfected with LRRK2 wt plasmid, with or without PMA/Ionomycin, and with 200 IU/mL IFN-γ for 24 hrs, where indicated. Data are expressed as fold over corresponding LRRK2 wt transfected cell line (mean ± SD, one-way ANOVA, Bonferroni post hoc, **P<0.01, *P<0.05; n=4). **(G)** NFAT luciferase assay in scrb and IQGAP1 KD HEK293 NFAT reporter cells transfected with LRRK2 G2019S plasmid, with or without PMA/Ionomycin, and with 200 IU/mL IFN-γ for 24 hrs, where indicated. Data are expressed as fold over corresponding LRRK2 G2019S transfected cells (mean ± SD, one-way ANOVA, Bonferroni post hoc, ***P<0.001, **P<0.01, *P<0.05; n=4).

### IFN-γ impairs neurite outgrowth in human iPSC-derived neurons

We next sought to investigate whether the reduction of NFAT3 nuclear localization observed in LRRK2 G2019S PD patient neurons has functional consequences relevant to human disease. Given that NFAT signaling is essential for axonal outgrowth in response to neurotrophins and netrins ^42^, and LRRK2 has been implicated in neurite outgrowth defects ^29, 43, 44^, we examined whether NFAT signaling could play a role in LRRK2-mediated defects of neurite outgrowth. To this end, three isogenic pairs of LRRK2 G2019S PD neurons and corresponding controls were treated with MCV1, a potent specific inhibitor of calcineurin-mediated NFAT activation, or IFN-γ, and neurite length was quantified by immunostaining for β-Tubulin III. NFAT inhibition, as well as IFN-γ treatment, led to a significant decrease of neurite length in both control and LRRK2 PD iPSC-derived neurons (Fig. 5A, B). The effect of IFN-γ on neurite length was confirmed by immunostaining for TH (Supplementary Fig. 2A, B). Notably, treatment with the LRRK2 kinase inhibitors IN-1 and GSK2578215A significantly rescued neurite length defects upon IFN-γ treatment (Supplementary Fig. 2C, D). To confirm that the reduction of neurite length was mediated by decreased NFAT activity, we performed a rescue experiment with Neuregulin 1 (NRG1) that promotes NFAT activity 45. Control and LRRK2 G2019S PD patient neurons were treated with MCV1 or IFN-γ with or without NRG1. Quantification of neurite length showed that NRG1 rescues neurite length defects in MCV1 and IFN-γ-treated control neurons (Fig. 5C, D). NRG1 treatment also partially rescued neurite length defects in LRRK2 G2019S neurons (Fig. 5C, D).

**Figure 5.**
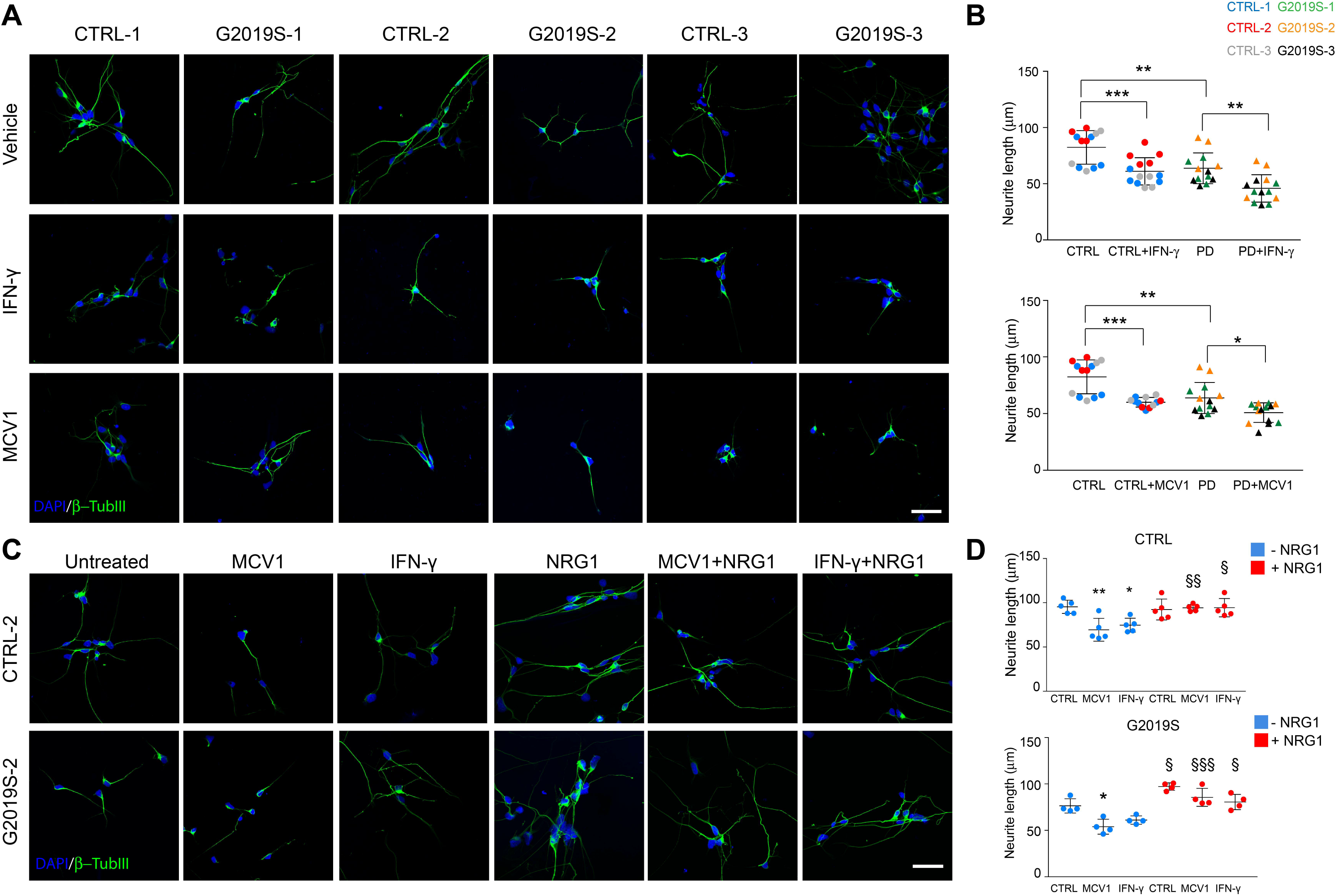
IFN-γ reduces neurite outgrowth by decreasing NFAT signaling. **(A)** Representative images of βTubIII immunostaining (green) showing neurite elongation in LRRK2 G2019S and isogenic control iPSC-derived neurons after treatment with vehicle, 500 nM MCV1 (NFAT inhibitor) for 24 hrs, or 200 IU/mL IFN-γ for 24 hrs. Blue, DAPI staining. Scale bars, 50 μm. **(B)** Quantification of neurite elongation is shown (mean ± SD, t-test, ***P<0.001, **P<0.01, *P<0.05; n=5 independent experiments). **(C)** Immunostaining for βTubIII (green) showing neurite elongation in LRRK2 G2019S and isogenic control iPSC-derived neurons after treatment with vehicle, 500 nM MCV1 24 hrs, or 200 IU/mL IFN-γ for 24 hrs with/without 200 ng/mL NRG1 treatment for 48 hrs. Scale bars, 50 μm. **(D)** Quantification of neurite elongation is shown (mean ± SD, one-way ANOVA, Bonferroni post hoc; for vehicle vs MCV1/IFN-γ treatment **P<0.01, *P<0.05; for NRG1 treatment §P<0.05, §§P<0.01, §§§ P<0.001; n=5 independent experiments).

### LRRK2 regulates NFAT shuttling in human macrophages and microglia

NFAT regulates responses within the innate and adaptive arms of the immune system ^46^. Since NFAT1 was the main isoform expressed in human microglia (Supplementary Fig. 1D), we focused on this isoform for subsequent analysis. We confirmed that IFN-γ increases LRRK2 expression in mononuclear phagocytes, by treating the monocytic cell line THP-1, differentiated to macrophages, and human iPSC-microglia with 200 IU/mL IFN-γ for 72 hours. Similar to what observed in neurons, LRRK2 expression was increased after IFN-γ stimulation (Supplementary Fig. 3A). To investigate the role of LRRK2 in NFAT1 shuttling in human macrophages, we knocked down LRRK2 (LRRK2 KD) via lentiviral-mediated shRNA delivery in THP-1 cells (Supplementary Fig. 3B, C). LRRK2 KD THP-1 macrophages displayed increased NFAT1 nuclear shuttling upon stimulation with ionomycin (Supplementary Fig. 3D, E). Control, LRRK2 KO iPSCs, as well as G2019S iPSCs efficiently differentiated into microglia without significant differences (Fig. 6A-C, (Supplementary Fig. 4A). Interestingly, microglia motility was increased in LRRK2 G2019S microglia compared to control (Fig. 6D). However, IFN-γ significantly decreased motility of G2019S, but not control, microglia (Fig. 6D). Immunostaining analysis of NFAT1 subcellular localization revealed increased levels of nuclear NFAT in LRRK2 KO microglia treated with ionomycin (Supplementary Fig. 4A, B). On the contrary, nuclear NFAT levels were significantly reduced in LRRK2 G2019S microglia (Supplementary Fig. 4A, B). We next examined the functional consequences of NFAT activity in human microglia. Given that ionomycin is a weak stimulator of cytokine release in microglia, we employed LPS stimulation to induce NFAT nuclear shuttling, based on previous reports ^47^. Nuclear NFAT1 was significantly increased in LRRK2 KO compared to CTRL microglia upon LPS treatment, whereas a significant reduction was observed in LPS-stimulated LRRK2 G2019S cells compared to isogenic controls (Supplementary Fig. 4A, C). Next, control, G2019S and LRRK2 KO iPSC-derived microglia were treated with LPS and cytokine production was assessed by Multiplex Elisa (Fig. 6E, Supplementary Fig. 5A, B). Among the analyzed cytokines assessed by Multiplex Elisa, IL-6, TNF-α, IL-8 significantly decreased in G2019S compared to isogenic CTRL microglia, whereas IL-10, IL-1β, IL-12p70, VEGF and MIP-1β were significantly increased, upon LPS stimulation (Fig. 6E, Supplementary Fig. 5A). On the other hand, we detected a significant increase of IL-6 and MIP-1β as well as a significant down-regulation of IL-1β and IL-10 secretion in LPS-treated LRRK2 KO microglia compared to isogenic wt controls (Fig. 6E). Next we further investigated the impact of LRRK2 on the metabolic and inflammatory profile of LPS-stimulated microglia. To this end, we examined the extracellular acidification rate (ECAR) by Seahorse metabolic analysis, which revealed a defect of the LPS-induced glycolytic switch in LRRK2 KO microglia as well as in control microglia upon treatment with the LRRK2 inhibitor IN-1 (Fig. 6F). To further confirm the impact of LRRK2 G2019S on the pro-inflammatory profile, we examined the response of microglia to other TLR agonists that elicit a type II IFN response. To this end, isogenic control, LRRK2 G2019S and LRRK2 KO microglia were treated with the viral mimic polyinosinic:polycytidylic acid (poly(I:C),10μg/mL) and production of IL-1β was assessed by Elisa. Levels of IL-1β were significanty decreased in poly(I:C)-stimulated LRRK2 KO microglia compared to CTRL microglia. Poly(I:C)-treated LRRK2 G2019S microglia showed an increase in IL-1β production compared to isogenic controls, although this increase was not significant (Fig. 6G). Finally, to study the effects of microglia on neurons, iPSC-derived neurons were exposed to conditioned media from LPS-activated microglia (microglial-conditioned media-MCM-). The findings showed that only MCM from LRRK2 G2019S activated microglia contributes to an inflammatory/toxic environment that affects neurite elongation (Fig. 6H and (Supplementary Fig. 5B).

**Figure 6.**
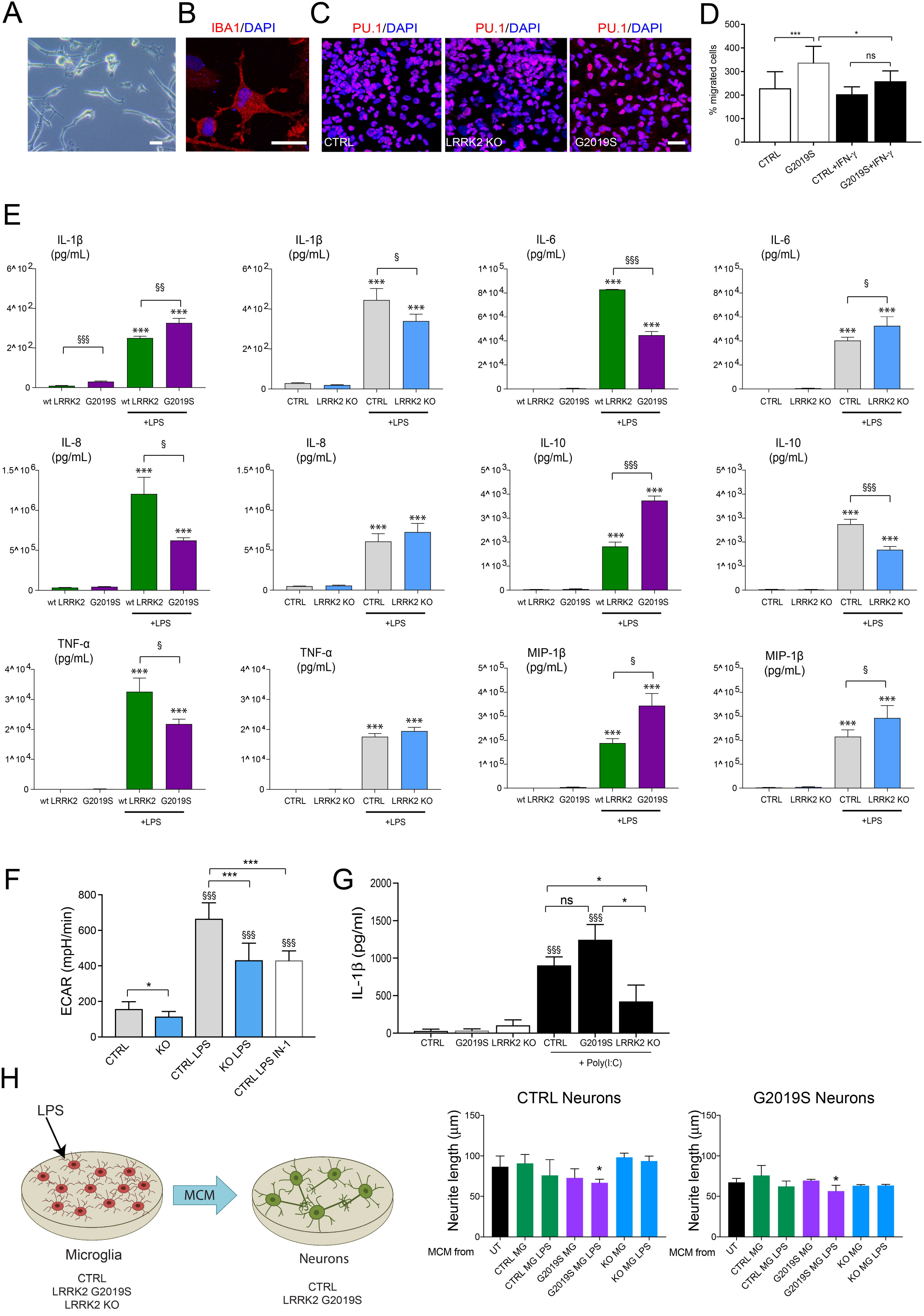
LRRK2 regulates NFAT activity and cytokine production in human iPSC-derived microglia. **(A)** Representative bright-field image of ramified iPSC-derived microglia. Scale bar, 20 μm. **(B)** Representative confocal image of iPSC-derived microglia immunostained for IBA1 (red) and nuclear staining DAPI (blue). Scale bar, 20 μm. **(C)** Representative confocal images of iPSC-derived microglia immunostained for PU.1 (red), and nuclear staining DAPI (blue). Scale bar, 20 μm. **(D)** Quantification of migrated iPSC-derived microglia cells upon 100 μM ATP stimulation, with or without 200 IU/mL IFN-γ treatment (mean ± SD, one-way ANOVA, Bonferroni post hoc; ***P<0.001, *P<0.05; n=3). **(E)** Isogenic CTRL, LRRK2 KO, and LRRK2 G2019S iPSC-derived microglia were treated with 100 ng/mL LPS for 24 hrs or left untreated, and cytokine levels were measured by Multiplex Elisa. Concentration of differentially secreted cytokines is shown (mean ± SD, t-test, treated vs untreated, ***P<0.001; G2019S vs CTRL or LRRK2 KO vs CTRL §P<0.05, §§P<0.01, §§§ P<0.001; n=3). **(F)** Seahorse metabolic analysis of extracellular acidification rate (ECAR) in CTRL and LRRK2 KO microglia treated with 100 ng/mL LPS with or without LRRK2 inhibitor 1.5 μ◻ IN-1 (mean ± SD, t-test, treated vs untreated, §§§ P<0.001; CTRL vs LRRK2 KO, CTRL LPS vs LRRK2 KO LPS or CTRL LPS IN-1, ***P<0.001, *P<0.05; n=8). **(G)** Isogenic wt (CTRL), LRRK2 KO, and LRRK2 G2019S iPSC-derived microglia were treated with 10 μg/mL poly(I:C) for 24 hrs and IL1-β levels were measured by Elisa (mean ± SD, t-test, treated vs untreated, §§§ P<0.001; CTRL treated vs G2019S treated or LRRK2 KO treated, *P<0.05; n=4). **(H)** Schematic diagram showing treatment of neurons with either untreated, or LPS-treated, microglial-conditioned medium (MCM); microglial cell lines used are indicated below the scheme. Quantification of neurite elongation in CTRL and G2019S neurons is shown on right (mean ± SD, t-test vs CTRL MG, *P<0.05; n=3 independent experiments).

## Discussion

Our study provides further understanding of the role of IFN type II signaling in PD and shows that LRRK2 G2019S mutation synergizes with IFN-γ, thus, serving as a potential direct link between inflammation and neurodegeneration in PD.

Besides their role in immune defense, both type I and type II IFNs contribute to physiological brain functions such as regulation of neurogenesis and synaptic function ^9, 48^. However, increasing evidence indicate that dysregulation of IFN signaling plays a role in ageing and neurodegenerative disease processes ^2, 3, 4^. With respect to PD, genetic and functional studies have identified a link between IFN-γ and disease ^5, 6, 11, 13, 14, 15^. There is also considerable evidence that PD genes can synergize with inflammatory triggers in determining disease risk ^16, 49, 50^. While PD genes can increase the vulnerability of nigral DA neurons to inflammation-related degeneration, they could also contribute to the dysregulation of immune responses to infections and the enhancement of age-related immune dysfunction ^51^. This connection is particularly relevant for LRRK2 mutations, which are the most frequent genetic cause of familial and sporadic PD ^17^. LRRK2 function goes beyond the brain and PD, being involved in inflammatory bowel diseases, infections, and cancer ^52, 53^. Consistent with a role for LRRK2 during infections, the *LRRK2* promoter region contains binding sites for IFN-response factors and IFN-γ robustly induces LRRK2 expression in a variety of immune cells ^18, 19, 20, 21, 22^. Using iPSC-based disease modeling, we have shown that IFN-γ induces LRRK2 expression also in human neurons. Thus, IFN-γ may serve as the central regulator of LRRK2-mediated responses during infections and immunity, as well as neuroinflammatory processes in a cell-type specific manner. Human neurons exhibited a strong IFN-γ response characterized by the upregulation of genes involved in cellular stress responses (*EIF2AK2*), apoptotic pathways (*IRF1, SOCS1*), brain development and axonal regeneration (*AKT3*, *SOCS1)*, and inflammasome assembly (*EIF2AK2*). However, IFN-γ alone did not elicit neuronal loss, in line with its primary role, which is to promote cellular integrity during infections ^54^. We found that IFN-γ reduces AKT phosphorylation in human neurons. Interestingly, IFN-γ treatment also suppresses AKT signaling in human macrophages, supporting the evidence that neuronal IFN-γ signaling is similar to that of immune cells ^55^. Previous work has implicated LRRK2 in AKT phosphorylation and shown that disease associated mutations reduce its interaction with and phosphorylation of AKT ^56, 57, 58^. In line with these findings, we found that IFN-γ synergizes with LRRK2 G2019S mutation to suppress AKT phosphorylation. Given the role of AKT in the regulation of NFAT activity ^31, 32^, we explored the impact of IFN-γ on NFAT signaling and its possible negative regulation of NFAT along with LRRK2 ^33^. In support of the study of Liu et al. ^33^, we found that LRRK2 is a negative regulator of NFAT activity in a variety of human cell types, including HEK293, human macrophages as well as iPSC-derived neurons and microglia. Furthermore, we have identified a role for IFN-γ as negative regulator of NFAT. This effect was mediated by the IFN-γ-dependent induction of LRRK2 levels that acts as a negative regulator of NFAT. On the other side, IFN-γ-dependent activation of the proteasome ^34, 35^ leads to degradation of NFAT. It is likely that the activation of the proteasome is mediated by the increased levels of SOCS1, one of the induced IFN-γ target genes, that activates the ubiquitin-proteasome pathway ^59^. Importantly, LRRK2 G2019S mutation results in even more pronounced effects on NFAT activity, due to its negative impact on AKT phosphorylation that in turn impairs NFAT translocation ^31, 32^.

In the original paper by Liu et al., LRRK2 has been shown to negatively modulate NFAT via NRON complex mediated cytosol retention ^33^. Using overexpression systems and PD patient cells, we have identified alternative mechanisms of this negative modulation and cell-type specific effects of reduced NFAT transcriptional activity. We show that LRRK2 G2019S negatively modulates NFAT shuttling by two distinct mechanisms: 1) by altering Ca^2+^ dynamics and 2) via a microtubule-dependent pathway. The main activation pathway of NFAT is the raise of intracellular Ca^2+^ via cell-surface receptors resulting in the activation of phospholipase C (PLC)-γ and Ca^2+^ release from ER via the IP3 receptor ^60^. Consistent with a recent report ^44^, we show that LRRK2 G2019S leads to increased intracellular Ca^2+^ levels at basal conditions and depletion of ER Ca^2+^ stores in PD patient neurons. However, LRRK2 G2019S neurons showed significantly reduced Ca^2+^ influx upon KCl depolarization. In agreement with the defect in Ca^2+^ signaling, LRRK2 G2019S neurons showed a marked decrease in the activation of NFAT. We have examined the impact of chronic IFN-γ treatment on intracellular Ca^2+^ dynamics. While it is known that acute IFN-γ treatment induces Ca^2+^ transients ^61, 62^, our data show that chronic IFN-γ leads to a reduction of intracellular Ca^2+^ levels and depletion of Ca^2+^ in the ER stores that could explain the reduced NFAT nuclear localization. Importantly, we found that IFN-γ synergizes with LRRK2 G2019S to enhance NFAT inhibition via Ca^2+^ mediated mechanisms.

Integrity of the microtubule network is necessary for NFAT translocation and T-cell activation ^36^. LRRK2 plays a physiological role in cytoskeletal organization and pathogenetic variants alter microtubule stability with a cell-type specific functional impact ^37^. Based on these premises, we hypothesized that IQGAP1, a component of the NRON complex involved in cytoskeleton organization, can be the connecting element between IF-γ, LRRK2 and dysfunctional NFAT shuttling. IQGAP1 contributes to numerous brain functions including neurogenesis, neurite outgrowth, and maintenance of neurons ^40^. Interestingly, IFN-γ leads to cytoskeleton reorganization that may be partially mediated by IQGAP1 ^63^. As hypothesized, our knockdown experiments show that IQGAP1 is involved in LRRK2 and IFN-γ-mediated negative regulation of NFAT shuttling.

Finally, we have identified cell-type specific effects of reduced NFAT translocation. Consistent with the role of calcineurin/NFAT signaling in the regulation of axon outgrowth ^42^, we show that NFAT signaling plays a role in neurite outgrowth in human neurons and that IFN-γ synergizes with LRRK2 G2019S to inhibit NFAT activity and determine a reduction of neurite length. Therefore, our results provide further mechanistic explanation of the neurite growth defects that have been observed in LRRK2 G2019S neurons. To examine the cell-specific role of reduced NFAT activity, we have employed human macrophages and human iPSC-derived microglia. Whether adult microglia express LRRK2 is still controversial; our data show that LRRK2 is expressed by human microglia at basal conditions and its expression is enhanced by IFN-γ. In addition, we show that human microglia produce IFN-γ when stimulated by LPS. However, we cannot exclude that the *in vitro* culture conditions used in the present study enhance LRRK2 expression and cytokine production. In line with its role in cytoskeletal remodeling ^63^, IFN-γ stimulation reduced the motility of microglia. However, such an effect was significant only in LRRK2 G2019S microglia, suggesting that PD LRRK2 mutations enhance immune responses to IFN-γ

Importantly, we show that LRRK2 deficiency in innate immune cells leads to increased NFAT nuclear localization. On the contrary, LRRK2 G2019S was associated to decreased nuclear NFAT. Upon stimulation with LPS, which is known to promote NFAT nuclear translocation in microglia ^47^, we detected LRRK2- and NFAT-dependent changes in cytokine production. Specifically, IL-6, TNF-α, IL-8, and MCP-1 production was decreased in G2019S microglia with opposite changes in LRRK2 KO microglia. However, G2019S microglia showed increased production of, IL-12p70 and MIP-1β upon stimulation, suggesting that additional mechanisms other than NFAT regulate cytokine responses in LRRK2 KO and G2019S microglia. In this respect, we show that LRRK2 modulates microglia activation by interfering with the metabolic switch toward glycolysis that normally occurs in LPS-activated macrophages ^64^. Importantly, activated G2019S microglia leads to neurite shortening, suggesting a neurotoxic effect of LRRK2-driven immunological changes in PD.

Increased IFN-γ levels have been documented in PD patients ^11, 12^. The source of IFN-γ in the aged and PD brain still needs to be identified. Our work shows that LRRK2 does not influence IFN-γ production in microglia, suggesting that microglial cells are not the primary source of IFN-γ. Ageing is accompanied by immune dysfunction and a paradoxical increase of cytokines in several tissues, including the brain ^51^. A recent study conducted in mice demonstrates a clonal T cell expansion within the brain of old mice with high levels of IFN-γ, suggesting a role for T cell-derived IFN-γ in the age-dependent decline of brain function ^65^. Given the presence of T lymphocytes in PD brains and the increasing evidence pointing towards a role for T cell responses in disease pathogenesis ^66, 67, 68^, T-cells could be the main source of IFN-γ, which, in turn, induces LRRK2 expression in neurons and microglia, leading to neuronal damage and inflammatory reactions. Thus, further studies are warranted to investigate the role of IFN signaling in specific brain regions and the role of IFN-γ producing T-cells in ageing and PD.

In summary, our work shows that LRRK2 G2019S sensitizes neurons to IFN-γ inflammatory responses, thus serving as a potential direct link between inflammation and neurodegeneration in PD. We have uncovered a cell type-specific role for the synergistic effect between IFN-γ and LRRK2 that may influence PD risk through specific ageing signatures that converge on the immune system. Importantly, given the role of LRRK2 in sporadic PD ^69, 70^, the relevance of our work goes beyond LRRK2 G2019S patients and suggests that the IFN-γ /LRRK2 axis may be a potential target for intervention in acute or chronic states of neuroinflammation in genetic and sporadic PD.

## Methods

### Differentiation of NPCs into midbrain DA neurons

All cells used in the study were derived from patients who signed an informed consent. The Ethics Committee of the Medical Faculty and the University Hospital Tübingen (Ethikkommission der Medizinischen Fakultät am Universitätsklinikum Tübingen) approved the protocol before performing the experiments. Patient LRRK2 G2019S and corresponding gene-corrected isogenic control iPSC-derived NPCs were previously generated using an NPC-based protocol ^28^. NPC differentiation into DA neurons was done according to a previously published protocol. NPCs were passaged with Accutase (Sigma-Aldrich) every 5 to 6 days on Matrigel-coated plates at 1:5-1:10 ratio in expansion medium containing N2/B27 medium, 150 μ ascorbic M CHIR 99021 (CHIR, Axon Medchem), and 0.5 μM purmorphamine (PMA, EMD). NPCs were plated for further differentiation and, when they reached 70% confluency, expansion medium was replaced by N2/B27 medium supplemented with 200 μ AA, 100 ng/mL FGF8 (Peprotech) and 1 μ PMA, changed every other day. On day 8, maturation medium containing N2/B27 medium with 20 ng/mL BDNF (Peprotech), 500 μ β (Peprotech), 20 ng/mL GDNF (Peprotech), 200 μ AA, and 0.5 μM PMA (for 2 more days) was applied to the cells. After switching to maturation medium, confluent cultures were detached with Accutase, and replated at a 1:3 ratio on Matrigel-coated plates. Cells were kept for 2 weeks in maturation medium, before performing experiments. Cell lines were routinely assessed by Sanger sequencing for cell line identity, as well as for mycoplasma (every 2 months). Immunostaining for β III Tubulin and TH was also performed to evaluate the quality of the neuronal differentiation. Only cultures that properly differentiated into dopaminergic neurons were used for experiments.

### Reagents

Where indicated, cells were treated with recombinant human IFN-γ (R&D), LPS (LPS from E. coli O111:B4) (Invivogen), polyinosinic:polycytidylic acid (Invivogen), Ionomycin (Sigma-Aldrich), LRRK2-IN-1 and GSK2578215A (Tocris).

### Generation of LRRK2 knock out iPSCs

LRRK2 KO iPSCs (clone #1) was generated using zinc finger nucleases ZFNs (Sigma). Cells were transfected with 2 μg of each ZFN construct, as well as 2 μg linearized targeting vector harboring a premature stop codon in LRRK2 exon 41 and a neomycin resistance cassette, using Amaxa Nucleofector II (Lonza), Nucleofection Solution for human stem cells II (Lonza), and program B-16, according to the manufacturer’s instructions. Cells were replated onto MEF feeder-coated plates in hESC medium supplemented with 10 μM ROCK inhibitor (Ascent Scientific). After colony formation, selection for homologous recombination was performed by adding 50 μg/mL G418 (PAA). Resistant colonies were picked and clonally expanded. A second step of nucleofection was performed with 2 μg of the same homologous construct as before, harboring a blasticidin resistance cassette. Selection was performed with 100 μg/mL of Blasticidin (InvivoGen), and resistant colonies were expanded on MEF feeder-coated plates. The presence of the stop codon in exon 41 was validated by Sanger sequencing (primers FW: gcacagaatttttgatgcttg; RV: gaggtcagtggttatccatcc). LRRK2 KO clones (clone #2 and #3) were generated from a newly generated viral free control iPSC line (CTRL-4). For iPSC generation, human fibroblasts were cultured in fibroblast culture medium [DMEM high glucose (Life Technologies) plus 10% FBS (Life Technologies)]. Reprogramming was performed by nucleofection with the episomal plasmids pCXLE-hUL, pCXLE-hSK and pCXLE-hOCT4 as described by Okita et al. ^71^. Pluripotency was assessed by immunostaining for pluripotency genes (OCT4, SOX2, TRA-1-60, TRA-1-80) and EB-based differentiation. To verify the absence of integration of the reprogramming plasmids, RT-PCR was performed with plasmid-specific primers ^71^. Genomic integrity of selected clones was assessed by G-Banding karyotyping and high-density genotyping using the Illumina HumanOmni2.5-8 (Omni2.5) array. Copy number analysis was performed using CNVPartition plugin (Illumina). Synthetic sgRNAs targeting LRRK2 exon 3 were purchased from Synthego (sgRNA1: CAA UCA UUU CCA ACA UCC UG; sgRNA2: AAA UUA AUA GAA GUC UGU CC). Nucleofection of 180 pmol sgRNA1, 180 pmol sgRNA2, 20 pmol Cas9 nuclease (sgRNA to Cas9 nuclease ratio used as 9:1) was performed with Amaxa 2D Nucleofector (Lonza, program B016). After recovery, iPSCs were clonally expanded and the genomic deletion assessed by PCR (FW: GGT GGG TTG GTC ACT TCT GT; RV: ACA GGA GTA GCC TTG GAT TGC) and Sanger sequencing (FW: ACG GCT TGC TTT TGT TTC TGG; RV: GCA AAC ACA GTG TAT CAA GGG A). The screening of possible off-target effects was performed using crispr.mit.edu and crispr.cos.uni-heidelberg.de; no exonic off-target effect was predicted. Genomic integrity of selected clones was assessed by G-Banding karyotyping and high-density genotyping using the Illumina HumanOmni2.5-8 (Omni2.5) array. iPSCs were kept in culture in hESC medium consisting of knockout DMEM (Gibco Life Technologies), 20% Knockout Serum replacement (Gibco Life Technologies), 1% penicillin/streptomycin (P/S, Merck Millipore), 1% non-essential amino acids (Gibco Life Technologies), 500 μM β-mercaptoethanol (Sigma-Aldrich), 1% GlutaMAX Supplement (Gibco Life Technologies), supplemented with 10 ng/mL FGF2 (Peprotech). iPSCs were maintained by splitting onto mitomycin-treated CF-1 mouse embryonic fibroblasts (MEF) (Globalstem MTI ThermoFisher) with medium μM Rock inhibitor Y-27632 2HCl (Selleckchem).

### Differentiation of human iPSCs into microglia

iPSCs were differentiated into microglia following an established protocol ^72^ with minor changes. In brief, embryoid bodies (EBs) were formed using AggreWells-800 (STEMCELL Technologies), and cultured in mTeSR1 (STEMCELL Technologies) plus bone morphogenetic protein 4 (BMP4, 50 ng/mL, Immunotools), vascular endothelial growth factor (VEGF, 50 ng/mL, Immunotools), and stem cell factor (SCF, 20 ng/mL, Immunotools), for four days with 75% medium exchange daily. On day four, EBs were collected and transferred into 6-well cell culture plate (12-16 EBs/well) in X-VIVO15 (Lonza) supplemented with IL-3, (25 ng/mL, Immunotools), macrophage colony-stimulating factor (M-CSF, 100 ng/mL, Immunotools), 2 mM Glutamax (Thermo Fisher), 1% penicillin/streptomycin (P/S, Merck Millipore) and 0.055 mM β l (Sigma Aldrich) with medium exchange weekly. After 0.055 mM β-mercaptoethano 3-4 weeks, floating cells were collected and seeded 100,000 cells/cm^2^ in matrigel-coated (Corning) cell culture plate in Advanced DMEM/F12 (LifeTechnologies) plus N2 supplement (Thermo Fisher), Glutamax, P/S, β l, M-CSF (100 ng/mL), IL-34 (100 ng/mL, Peprotech) and GM-CSF (10 ng/mL, Immunotools) with medium exchange twice a week.

### Experiments with microglia-conditioned medium (MCM)

The conditioned medium was collected after 24 hours from control microglia or 100 ng/mL LPS-activated microglia. MCM was then filtered, diluted 1:1 with neuronal medium and applied to neurons. Cells were fixed after 48 hrs and used for neurite length assay.

### THP-1 experiments

Human THP-1 cell lines were purchased from Sigma Aldrich and cultured in RPMI 1640 (Life technologies), 2mM Glutamax, and 10% FBS. THP-1 cells were differentiated into macrophages with 25 ng/mL phorbol 12-myristate 13-acetate (PMA, Sigma) for 48 hrs. For LRRK2 knockdown experiments, the following high titer lentiviruses were purchased from Sigma Mission Library: TRCN0000021462 10^8 TU vector pLK0.1, TRCN0000021460 10^8 TU vector pLK0.1, and SHC002V non-target control. Lentiviral infection of THP-1 macrophages was performed with spinfection at MOI 10.

### Plasmids

The plasmids SF-tagged LRRK2 wt and SF-tagged LRRK2 G2019S were a kind gift from CJ Gloeckner. pEGFP-C1 NFAT3 was purchased from Addgene (Cat. No.10961).

### Luciferase assay

NFAT Reporter - HEK293 Cell Line (PKC/ Ca^2+^ Pathway) cell lines were purchased from Tebu-Bio and cultured in Dulbecco’s MEM (Merck) supplemented with 10% (vol/vol) FBS (Gibco) and 400 ug/mL of Geneticin (Life technologies). For HEK293 IQGAP1 knockdown experiments, the following high titer lentiviruses were purchased from Sigma Mission Library: TRCN0000047487 10^8 TU vector pLK0.1, TRCN0000047486 10^8 TU vector pLK0.1 and SHC002V non-target control. Lentiviral infection of HEK293 was performed at MOI 1. Where specified, cells were transfected with 8 μ of plasmid DNA (SF-tagged LRRK2 WT, or SF-tagged LRRK2 G2019S) per 10cm^2^ dish using polyethyleneimine (Polysciences) solution. After 48 hrs, cells were stimulated with phorbol-12-myristate-13-acetate (PMA, Sigma) 40 ng/mL and ionomycin (Sigma) 1 uM for 24 hours. Luciferase activity was measured with the Dual-Luciferase Reporter Assay System (Promega).

### Western blot

Lysates for whole cell protein analysis were obtained by resuspending cells with Tris-buffered Saline (TBS) supplemented with 0.05% NP40 with the addition of phosphatase and protease inhibitors (Roche) on ice, followed by centrifugation at 4°C for 30 minutes at 14.000 rpm. Where indicated, cells were treated with 200 IU/mL IFN-γ and 20nM MG132 (Sigma-Aldrich) for 24 hrs. After determining the protein concentration using BCA assay (Pierce), 50 −100 μg of protein were loaded on a 7%-12% polyacrylamide gel and then transferred on a methanol activated PVDF membrane (Millipore). Blot membranes were blocked for one hr with 5% milk or 5% BSA in TBS with 0.1% Tween 20 (TBST). Blots were then incubated overnight at 4 °C with primary antibody of interest diluted in blocking solution (Western Blotting Reagent, Roche). The appropriate species of HRP-conjugated secondary antibody (from Sigma-Aldrich) resuspended in 5% milk or 5% BSA was then applied to the membrane for one hour at room temperature. Amersham ECL Western Blotting Detection Reagent and Amersham Hyperfilm (both GE healthcare) were used to visualize the proteins. ImageJ software (version 1.52a) was used for densitometric analysis. Nuclear cytosolic separation was performed using NE/PER Nuclear and cytosolic extraction kit (ThermoFisher) according to manufacturer’s instructions. In brief, 2*10^6 cells were harvested with Accutase and washed with cold PBS. The cell pellet was resuspended in CER I buffer supplemented with phosphatase and protease inhibitors (Roche) to isolate cytosolic fraction and lysed using CER II buffer. Cells were centrifuged for 10 min at 12000 g and resulting supernatant contained cytosolic fraction. The cell pellet was washed twice with cold PBS to remove remaining cytosolic proteins. The pellet was resuspended in NER buffer supplemented with phosphatase and protease inhibitors (Roche) and repeatedly vortexed on ice for 40 minutes. 10 μg of protein was loaded on Tris-HCL Nupage gradient gel and the membrane was probed for cytoplasmic and nuclear compartment markers. Primary antibodies: rat monoclonal anti-LRRK2 24D8 (kind gift from C. Johannes Gloeckner; 2 μ rat monoclonal anti-LRRK2 1E11 (kind gift from C. Johannes Gloeckner; 4 μg); rabbit anti-AKT3 (1:1000, Cell Signaling Technology, Cat. No. 4059); rabbit anti-NFAT1 (D43B1) XP (1:1000, Cell Signaling Technology, Cat. No. 5861); rabbit anti-NFAT3 (23E6) (1:1000, Cell Signaling Technology, Cat. No. 2183); rabbit anti-PARP1 (1:8000, Cell Signaling Technology, Cat. No. 9542); rabbit anti HSP90 (1:20000, Enzo, Cat. No.ADI-SPA-836F); mouse anti-β-Actin (1:20000, Sigma, Cat. No. A5441); rabbit anti-IQGAP1 (1:500, Cell Signaling Technology, Cat. No. 2293); rabbit anti-AKT (1:1000, Cell Signaling Technology, Cat. No.9272); rabbit anti-phospho-AKT (Ser473) (1:1000, Cell Signaling Technology, Cat. No.4060).

### Immunocytochemistry

Cells were fixed in 4% paraformaldehyde (PFA) in PBS (w/v) for 15 minutes, then washed with PBS and blocked with PBST (PBS with 0.1% TritonX-100) with 10% normal goat serum (NGS) for 30 minutes. Then, cells were incubated overnight at 4 °C with primary antibody of interest diluted in blocking solution. The appropriate species of Alexa Fluor488/568/647-conjugated secondary antibody (from Invitrogen) resuspended in PBST with 1% NGS was then applied to the cells for 1 hr at room temperature. Cell nuclei were counterstained with DAPI (Biozol), and images were acquired with a Zeiss Imager Z1 with Apotome (Carl Zeiss) microscope. For neurite length analysis, neurons were plate at low density (25,000 cells/cm^2^) and fixed for immunostaining after 72 hrs. Images were analyzed using the Fiji plugin “Simple Neurite Tracer”. Primary antibodies: Rabbit anti NFAT1 (D43B1) (1:1000, Cell Signaling Technology, #5861); mouse anti-Tubulin β3 (TUBB3) (1:1000, Covance, #MMS-435P); mouse anti-Anti-β-Tubulin (1:1000, Sigma, #T8328); rabbit anti-Tyrosine Hydroxylase (1:500, Pel-Freeze, #P40101-150); rabbit anti-DYKDDDDK Tag (D6W5B) (1:1000, Cell Signaling Technology, #14793S); mouse anti Spi1/PU.1 (1:100, Biolegend, #658002); rabbit anti-Iba1 (1:2000, Wako Chemicals, 016-20001). For NRG1 experiments, neuronal cultures were treated with 200 ng/mL recombinant human NRG1 (BioLegend, Cat. No. 551904) for 48 hrs. For NFAT inhibition experiments, cells were treated with 500 nM MCV1 (HPVIVIT, Calbiochem Cat. No. 480404) for 24 hrs.

### Cytokine analysis

Levels of IL-1β IL-1Ra, IL-4, IL-6, IL-8, IL-10, IL-12p70, IFN-γ, MCP-1, MIP-1β, TNFα and VEGF were determined using a set of “in house developed” Luminex-α based sandwich immunoassays, each consisting of commercially available capture and detection antibodies and calibrator proteins. Samples were diluted at least 1:4 or higher to receive results below the upper limit of quantification. After incubation of the pre-diluted samples or calibrator protein with the capture-coated microspheres, beads were washed and incubated with biotinylated detection antibodies. Streptavidin-phycoerythrin was added after an additional washing step for visualization. For control purposes, calibrators and quality control samples were included on each microtiter plate. All measurements were performed on a Luminex FlexMap® 3D analyzer system, using Luminex xPONENT® 4.2 software (Luminex, Austin, TX, USA). For data analysis MasterPlex QT, version 5.0 was employed. IL-1β levels after poly(I:C) stimulation were measured using Human IL-1 beta/IL-1F2 DuoSet ELISA (R&D).

### Quantitative RT-PCR

mRNA isolation and reverse transcription reaction were performed respectively with RNeasy Mini kit and with QuantiTect Reverse Transcription kit (both from Qiagen), according to manufacturer’s instructions. QuantiTect SYBR GREEN I kit (Qiagen) and a Viia7 Real time PCR system (Applied Biosystems) were used for quantitative PCR reaction. The expression level of each gene was normalized to the levels of the housekeeping genes ribosomal protein large P0 (Rplp0), Hydroxymethylbilane Synthase (HMBS), or β2-microglobulin. The 2-DDCT method was used to calculate fold-changes in gene expression, based on housekeeping genes and biological reference samples for normalization.

### Seahorse XF^e^96 Metabolic Flux Analysis

Extracellular acidification rate was analyzed using an XF^e^96 Extracellular Flux Analyzer (Seahorse Biosciences). iPSC-derived microglia were grown on V3-PS XF plates (Seahorse Biosciences) at a density of 70,000 per well for 14 days. Measurement of extracellular acidification rate was performed using Seahorse XFp analyzer (Agilent) in freshly prepared medium, consisting of phenol-free DMEM supplemented with 1mM Glutamine and pH adjusted to 7.4. Glycolytic function was evaluated after subsequent injection of 10mM Glucose, 10 μ Oligomycin and 50mM 2-Deoxy-D-Glucose (all Sigma-Aldrich); for each condition 3 measurements lasting 5 minutes each were performed. After measurement, values were normalized to cell number by counting DAPI stained nuclei using a high-content cell analyzer (BD Bioscience, Pathway 855).

### Microglial cell motility assay

Microglia were plated into the top of trans-well migration chamber (5 μm-pore polycarbonate filters in 24-wells; Corning) containing Adenosine triphosphate (ATP, 100 μM; Sigma), in the bottom chamber, to induce chemotaxis. Wells containing medium without ATP were used as control. After 4 hrs, cells were fixed with PFA (4%) for 15 min at room temperature. Cells on the upper side of the filter were wiped off using a cotton-swab. Cells were counted on the bottom side of the chamber under a light microscope. The number of migrated cells was expressed as % of the non ATP-stimulated, control condition.

### Calcium live-cell imaging

Neuronal cultures were loaded for 40 minutes with Fura-2 in a CO2 incubator at 37 °C (incubation solution: Bessel-medium with Fura-2-AM (0.27 μM) and Pluronic F127 (1:1 ratio), Thermo Fisher Scientific). The ratiometric recordings were carried out by an upright fluorescence microscope (BX50WI, Olympus) equipped with a 20X water immersion objective (LUMPlan FL, 20X/0.80W, ∞ /0, Olympus), a polychromator (VisiChrome, Till Photonics) and a CCD camera (RETIGA-R1, 1360×1024 pixels, 16 bit). During the recording, stacks (single-plane two-channel) of the cellular Fura-2 fluorescence at the focal plane were acquired at 2 Hz (λexc= 340 and 380 nm; Olympus U-MNU filter set, 40 ms exposure time, 8-pixel binning) using the VisiView software (Till Photonics). The cellular Ca^2+^-activity was recorded for 15 minutes, consisting of 5 minutes control-recording and following 10 minutes of post-recording under different conditions by application of 20 mM KCl (Sigma-Aldrich), 500 nM of Thapsigargin (Sigma-Aldrich) for 15 min of recording time or 200 IU/mL IFN-γ. Post-agent application, the active time of KCl was 1 minute followed by 3 minutes of washout (2 mL/min), while the other agents were not washed out during the post-recording. The reagent solutions (200 μL) were injected (pressure application via micro pipette) within the recording chamber solution in the vicinity of the objective, reaching the cells immediately. The temperature of the recording chamber was set to 37 °C. For data analysis, ratio-stacks of the Ca^2+^-imaging recordings were generated by dividing the fluorescence images recorded at the excitation wavelengths of F340 and F380. To detect cellular Ca^2+^-values, selected cells were manually encircled by regions of interest (ROIs) and the obtained ROIs coordinates were used to extract corresponding Ca2+ traces from the ratio-stacks (custom written ImageJ macros (https://imagej.nih.gov/ij/)). The data were scaled ([0 1]) for further analysis (y=(x-min(min(x)))/(max(max(x))-min(min(x))), processed by custom written Matlab scripts (Mathworks, USA)). The baseline-value was generated as the mean of 5 values taken at each minute, during the 5-minute control-recording time. The agent dependent increase of cellular Ca^2+^ levels was calculated as delta of peak-Ca^2+^ value post application and the pre-defined baseline value. The recovery-time of the KCl induced Ca^2+^ increase was acquired at ±5% of the baseline.

### Statistical analysis

The Statistical Package GraphPad Prism version 7.00 (GraphPad Software, San Diego California USA) was used to analyse the data. Statistical testing involved two-tailed Student’s t-test or One-way Anova followed by Bonferroni multiple comparison test, as indicated. Data are expressed as mean + SEM (or + SD) as indicated. Significance was considered for P<0.05.

## Acknowledgements

We thank Gabriele Di Napoli for excellent technical support. We also thank Christian Johannes Gloeckner for sharing with us the LRRK2 plasmids. This work was supported by German Center for Neurodegenerative Disease (DZNE) and the Helmholtz Association Young Investigator Award (VH-NG-1123, to M.D.); the Marie Curie Career Integration Grant MC CIG304108 (M.D.), DFG research grant (DE 2157/2-1), COEN grant 4008 (M.D.); Hector Foundation (W.H.), and Tistou and Charlotte Kerstan Foundation (W.H.).

## Author Contributions

M.D., S.D.C., D.I., V.P. conceived and designed the experiments. S.D.C., D.I., and V.P. performed most of the experiments and analysed data; W.H. and D.I. designed, performed, and analysed data relative to Ca^2+^ measurements; C.Y. and D.S. generated and characterized LRRK2 KO iPSCs; M.J.P performed experiments and analysed data; I.N. and A.A. designed and performed imaging experiments; D.S. performed initial experiments on interferon signaling; R.P.C. performed IQGAP1 experiments; N.S. and M.J. performed Multiple Elisa experiments; T.G. provided patient samples; M.D. supervised the study, acquired funding, and wrote the manuscript with contribution from all authors.

